# Mutations in the *SmAPRR2* transcription factor suppressing chlorophyll pigmentation in the eggplant fruit peel are key drivers of a diversified colour palette

**DOI:** 10.1101/2022.08.23.504925

**Authors:** Andrea Arrones, Giulio Mangino, David Alonso, Mariola Plazas, Jaime Prohens, Ezio Portis, Lorenzo Barchi, Giovanni Giuliano, Santiago Vilanova, Pietro Gramazio

## Abstract

Understanding the mechanisms by which chlorophylls are synthesized in the eggplant (*Solanum melongena*) fruit peel is of great relevance for eggplant breeding. A multi-parent advanced generation inter-cross (MAGIC) population and a germplasm collection have been screened for green pigmentation in the fruit peel and used to identify candidate genes for this trait. A genome-wide association study (GWAS) performed with 420 MAGIC individuals revealed a major association on chromosome 8 close to a gene similar to *APRR2*. Two variants in *SmAPRR2*, predicted as having a high impact effect, were associated with the absence of fruit chlorophyll pigmentation in the MAGIC population, and a large deletion of 5.27 kb was found in two reference genomes of accessions without chlorophyll in the fruit peel. The validation of the candidate gene *SmAPRR2* was performed by its sequencing in a set of MAGIC individuals and through its de novo assembly in 277 accessions from the G2P-SOL eggplant core collection. Two additional mutations in *SmAPRR2* associated with the lack of chlorophyll were identified in the core collection set. The phylogenetic analysis of *APRR2* reveals orthology within Solanaceae and suggests that specialization of *APRR2-like* genes occurred independently in Cucurbitaceae and Solanaceae. A strong geographical differentiation was observed in the frequency of predominant mutations in *SmAPRR2*, resulting in a lack of fruit chlorophyll pigmentation and suggesting that this phenotype may have arisen and been selected independently several times. This study represents the first identification of a major gene for fruit chlorophyll pigmentation in the eggplant fruit.

## Introduction

Chlorophylls are the most widely distributed and important natural pigments because of their essential role in photosynthesis (Gross, 2012; Jia et al., 2020). They are tetrapyrrole compounds, as are two other important plant cofactors, heme and phytochrome (Senge et al., 2014). Chlorophylls are also important photosensitizers, producing Reactive Oxygen Species (ROS) under saturating light intensities (Foyer, 2018). Their metabolism impacts the assembly of photosynthetic machineries but also influences processes such as programmed cell death, the ‘stay-green’ phenomenon, and chloroplast–nucleus communication (Tanaka and Tanaka, 2006). To optimize photosynthesis and cope with alterations in the light environment, plants have evolved a complex and highly regulated process of chloroplast development (Dubreuil et al., 2018). Given the increasing demand for plant products, as important food sources in the human diet and for many other industrial uses (Pérez-Gálvez et al., 2020), photosynthesis and chloroplast biogenesis have received intensive investigation for their positive involvement in crop nutritional quality (Llorente et al., 2020).

Leaves are the major organs of photosynthesis for most plants, where light energy is absorbed and received by chlorophyll molecules (Calvo et al., 2017). Green fruits also contain functional chloroplasts with an important photosynthetic activity that affects the fruit growth, development, and composition, leading to the accumulation of metabolites associated with nutritional quality (Calvo et al., 2017; Jia et al., 2020). Therefore, enhancing fruit chloroplast activity and accumulation can result in mature fleshy fruit with higher nutritional values (Powell et al., 2012; Pan et al., 2013; Jia et al., 2020).

Eggplant (*Solanum melongena* L.) is a widely grown crop, ranking third in global production among Solanaceae crops (Faostat, 2020). It is characterized by a high diversity of commercial fruit colours, which depend mainly on the presence or absence of two pigments: anthocyanins and chlorophylls (Daunay et al., 2004; Page et al., 2019). In eggplant, anthocyanins are responsible for the purple colour of fruit peel, one of the traits of greatest interest in eggplant breeding (Daunay and Hazra, 2012). Chlorophylls confer green colour to fruit peel that is visible when anthocyanins are absent or present in small concentrations. If anthocyanins are present, chlorophylls contribute to a darker background and reinforce the fruit darkness (Daunay et al., 2004). Purple-coloured eggplants are the most demanded in many markets (Li et al., 2018a), and developing dark purple-coloured eggplants, which result from the combination of anthocyanins with chlorophylls, is a major objective in eggplant breeding programs.

Candidate genes have been proposed for fruit anthocyanin synthesis, distribution, and accumulation (Moglia et al., 2020; Toppino et al., 2020; Mangino et al., 2022; Yang et al., 2022). However, causative genes for the presence of fruit chlorophylls in the eggplant fruit peel have not been elucidated yet. Although there are already some studies of QTLs in eggplant, none of them is focused on this trait. Up to now, it has only been observed that green fruit peel colour is dominant over the white one and the monogenic dominance of the green flesh over the white one, suggesting the dominance of chlorophyll presence over its absence (Daunay et al., 2004). Other studies reported that wild eggplant populations mainly exhibit fruits with green or greenish colour, whereas the white ones are more typical of some cultivated and weedy eggplants from southern India, conferring them the “egg-plant” or “vegetable egg” common name (Davidar et al., 2015; Mutegi et al., 2015). A single major QTL at the top of chromosome 8 was reported for a related trait, the presence of a green ring in the flesh next to the skin, and a member of the ferredoxin gene family was suggested as the best candidate due to its involvement in chlorophyll production (Portis et al., 2014).

The presence of chlorophylls in the eggplant fruit peel, not only for their influence on fruit colour but also for their potential effect on fruit composition, makes their study interesting for eggplant breeding. In this respect, the use of experimental and germplasm populations has been of great relevance for mapping quantitative trait loci (QTLs) and identifying candidate genes related to agronomic traits of interest (Gramazio et al., 2018). Here, for the first time, we identify a major candidate gene controlling the fruit skin chlorophyll biosynthesis in eggplant by using a multi-parent advanced generation inter-cross (MAGIC) population and a core germplasm collection for gene validation.

## Results

### Phenotypic variation and association analysis

MAGIC population founders are contrasting for fruit peel chlorophyll (FC), showing six out of eight founders the presence of FC, namely MM1597, DH ECAVI, MM577, AN-S-26, H15, and ASI-S-1 (founders A, B, C, D, E, and H, respectively), and two of them, A0416 and IVIA-371 (founders F and G, respectively), the absence of FC (Figure 1). The screening for FC segregation among the 420 S3MEGGIC individuals revealed considerable variation for this trait. The dominance of the FC presence was confirmed by the phenotype of the simple hybrids obtained by crossing founders with and without FC for developing the S3MEGGIC population (ExF and GxH hybrids) (Figure 1). No maternal effect was observed on FC in hybrids.

**Figure 1.**
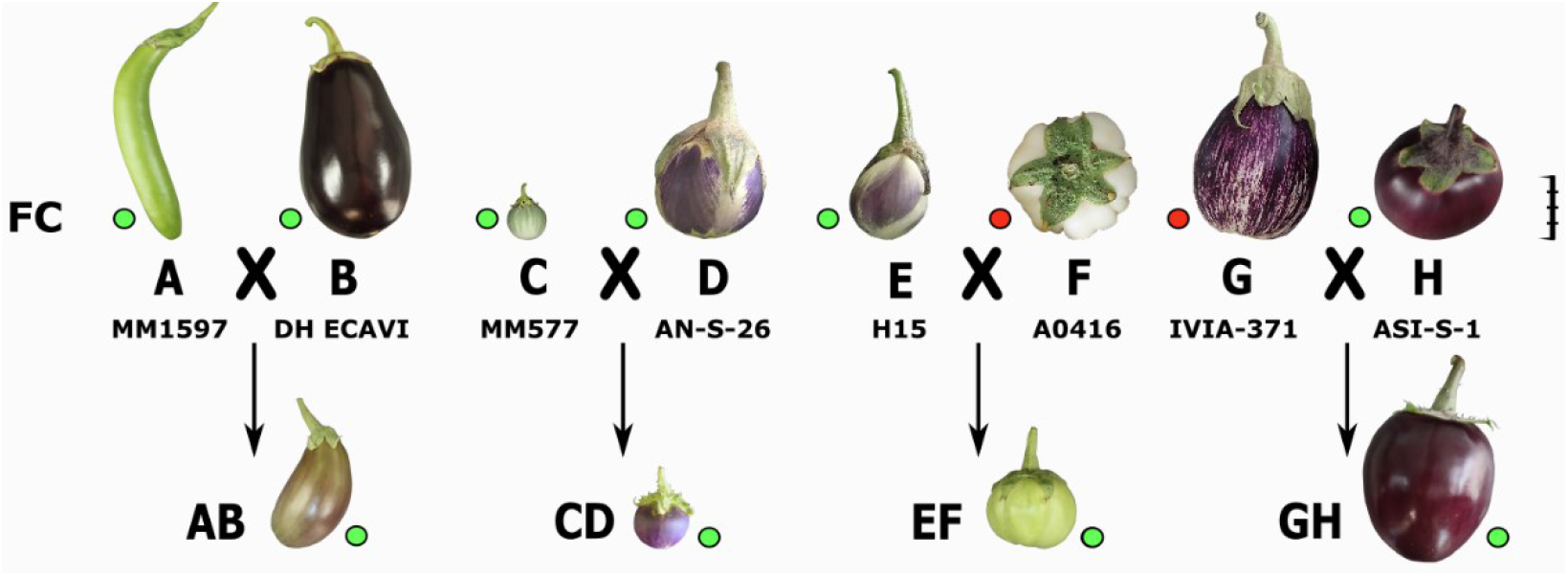
The eight founders, coded from A to H, from the S3MEGGIC population and the four simple hybrids obtained from their inter-cross (AB, CD, EF and GH) are represented at a scale based on the real fruit size. The scale bar represents 5 cm. Phenotyping for FC is represented by green and red dots, respectively.

GWAS analysis combining the genotypic and phenotypic data of the S3MEGGIC population led to the identification of significant associations for FC presence. The Manhattan plot revealed one major peak on chromosome 8, plus another one with a lower significance value on chromosome 4 (Figure 2). For the major peak on chromosome 8, 12 significant SNPs over the FDR threshold (LOD > 3.45) were identified, eight of them being over the Bonferroni threshold (LOD > 5.16) in the genomic region between 103.22 and 104.98 Mb and reaching the LOD of 34.88. On chromosome 4 there were eight significant SNPs over the FDR threshold (LOD > 3.45) in the region between 3.23 and 6.35 Mb, but none of them was significant over the Bonferroni threshold.

**Figure 2.**
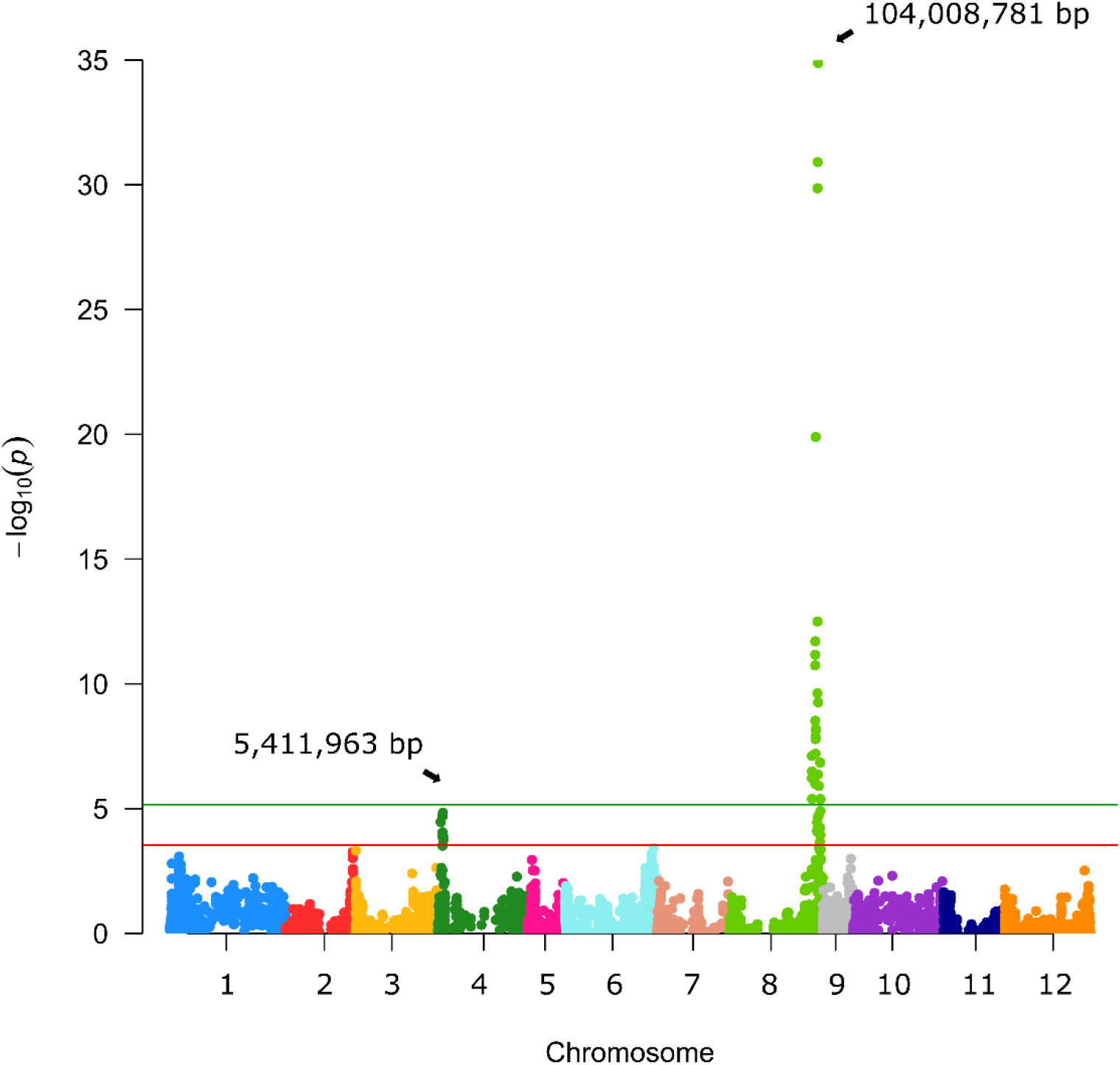
Manhattan plot for fruit chlorophyll. Arrows indicate the genome position of the highest peaks detected for the fruit peel chlorophyll trait. The red and green horizontal lines represent, respectively, FDR and Bonferroni significance thresholds at p=0.05.

For the main candidate region on chromosome 8, a comparative analysis of the founder haplotype diversity was performed in the S3MEGGIC population (Figure 3). The haplotype prediction of the 156 S3 lines with the lack of FC (i.e., having A0416 or IVIA-371 founder haplotypes) was successful in 91.03% of the cases. In this way, 50.00% of the cases (78 of 156) corresponded to the A0416 haplotype and 41.03% (64 of 156) to the IVIA-371 haplotype. In 6.41% of cases (10 of 156), the absence of FC was associated with the wild *S. incanum* MM577 founder haplotype, which presents some degree of heterozygosity that could have interfered with the association analysis. The remaining 2.46% (4 of 156) was evenly distributed between MM1597 and ASI-S-1 haplotypes, which could be the result of inaccurate phenotyping due to the presence of anthocyanins or of a small stylar scar, making phenotyping difficult.

**Figure 3.**
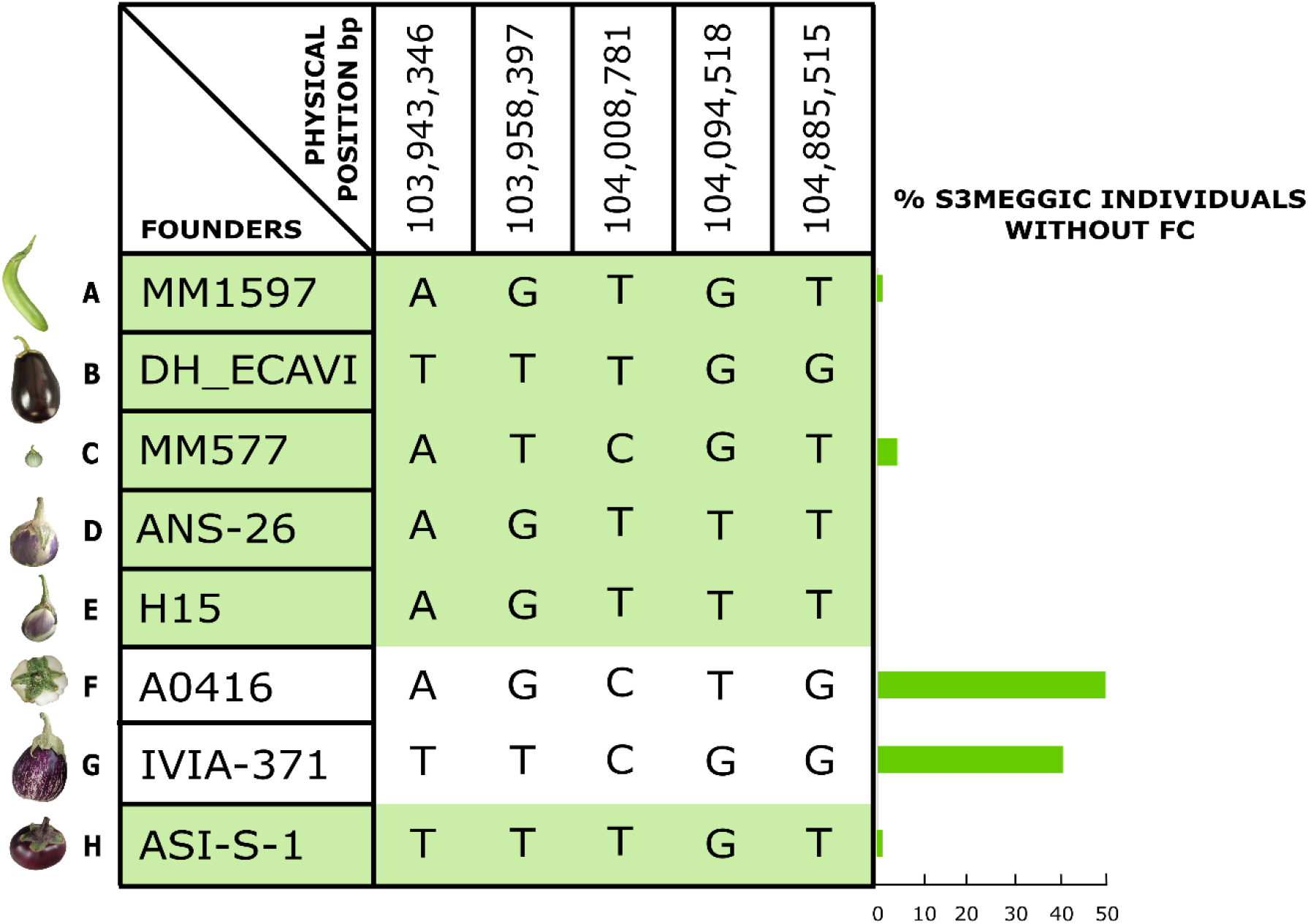
On the left, S3MEGGIC founder haplotypes at the most significant genomic region of chromosome 8; on the right, a histogram representing the percentage of S3MEGGIC individuals without FC with the same founder haplotype in that region.

### Candidate genes for chlorophyll biosynthesis

Based on the results of the GWAS analysis, putative candidate genes were identified close to or within LD blocks defined in the genomic regions with significant associations (Figure S1). From the S3MEGGIC founders’ resequencing data, variants with high impact effects on protein structure and function were annotated by SnpEff for all the candidate genes. Under the major GWAS peak of chromosome 8, in the genomic region of 104,008,667 – 104,012,138 bp, a candidate gene was identified as similar to ARABIDOPSIS PSEUDO RESPONSE REGULATOR2 (*APRR2*, SMEL_008g315370.1), which has been described as a pigment accumulation regulator and chloroplast development promotor in several solanaceous and cucurbitaceous crops (Pan et al., 2013; Liu et al., 2016; Oren et al., 2019). This gene presented high impact variants in those S3MEGGIC founders that do not present FC (founders F and G), which was confirmed by aligning all founder gene sequences retrieved and visualized in IGV. Specifically, founders F and G exhibited small deletions predicted to cause a frameshift variant identified as high impact: TG instead of TCTCCG in the 14,010,718 bp position on exon 3 and AT instead of ACT in the 104,011,578 bp position on exon 6, respectively (Figure 4a).

**Figure 4.**
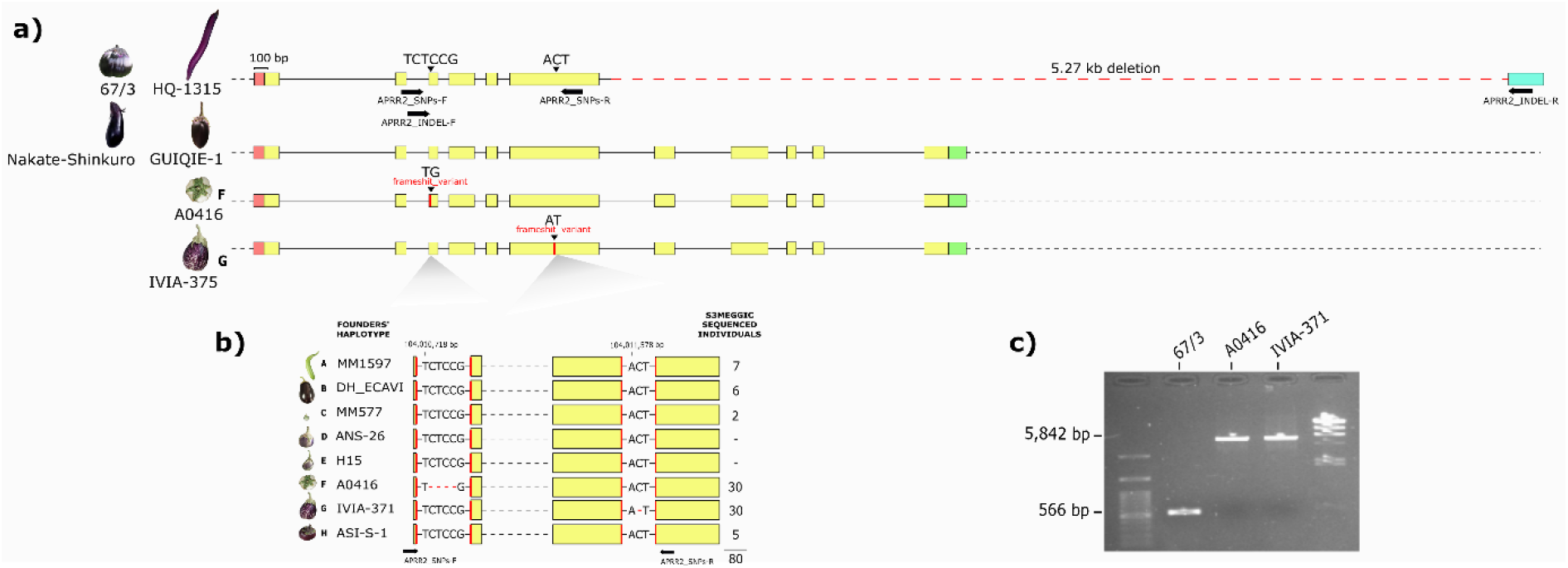
(a) *SmAPRR2* gene structure (5’-UTR in red, exons in yellow, 3’-UTR in green and 3’-UTR annotated in the 67/3 eggplant reference genome in blue) for 67/3, HQ-1315, Nakate-Shinkuro, GUIQIE-1 reference genomes and A0416 and IVIA-375 (S3MEGGIC founders F and G, respectively). Structural variations are indicated with black arrowheads and primers hybridization positions with horizontal black arrows. (b) The 80 S3MEGGIC selected individuals for sequencing based on their haplotype and their respective *SmAPRR2* gene sequence between APRR2_SNPs primers. (c) Electrophoresis gel of the PCR amplification using APRR2_INDEL primers for 67/3 and S3MEGGIC founders F and G genomes.

Same procedure was performed for the minor peak on chromosome 4. All the candidate genes close to or within the LD block including the genomic region with significant association were annotated by SnpEff for each of the S3MEGGIC founders. In the genomic region of 5,457,658 – 5,461,306 bp, a candidate gene was identified as similar to *GLK2* (SMEL_004g203570.1), which has been described as a positive regulator of chloroplast development and pigment accumulation in tomato fruit (Powell et al., 2012; Nguyen et al., 2014). Although according to the *SmGLK2* annotation it might fit with the regulation of chlorophyll biosynthesis, no high-effect variants were predicted by SnpEff in this gene for any of the founders without FC.

### Variants on SmAPRR2 sequence

To confirm experimentally the high impact variants predicted by SnpEff for S3MEGGIC founders F and G in the *SmAPRR2* gene sequence, 80 S3MEGGIC individuals were selected based on their phenotype and haplotype (20 individuals with FC, 30 with A0416 haplotype and 30 with IVIA-371 haplotype). The *SmAPRR2* gene sequence of the selected individuals was amplified by PCR using the primers combination APRR2_SNPs, which amplified a region of 1,363 bp where the two high impact variants were located (Figure 4a, Table S1). The Sanger sequencing of the amplicon confirmed the presence of specific allelic variants according to their phenotype (Figure 4b). The T(CTCC)G deletion was found in each of the 30 individuals with A0416 haplotype and the A(C)T deletion was found in each of the 30 individuals with IVIA-371 haplotype. As expected, no other variants were identified in the 20 remaining individuals that presented FC.

To further investigate the *SmAPRR2* gene structure, its nucleotide sequence was retrieved from the reference genome 67/3 (ver. 3, Barchi et al., 2019b), which corresponds to an accession without FC (SMEL_008g315370.1), presenting a 3,471-bp gene comprising 6 exons. The same gene structure was observed also in the HQ-1315 eggplant reference genome (Wei et al., 2020), which corresponds to another accession without FC (Smechr0802018). However, in the Nakate-Shinkuro (Sme2.5_00446.1_g00003.1) and the GUIQIE-1 eggplant reference genomes (Hirakawa et al., 2014; Li et al., 2021), both of which exhibit FC, a larger gene size of 4.794 bp with 11 exons was found (Figure 4a). Comparing the four reference genomes for the *SmAPRR2* gene structure, a large deletion of 5.27 kb was identified on 67/3 and HQ-1315 accessions causing the loss of exons 7 to 11. This large deletion did not correspond with the two high-effect variants observed in the S3MEGGIC population, where the larger gene size of 11 exons was the only *SmAPRR2* gene structure present in the population. The presence of the large deletion in the 67/3 accession was confirmed by PCR, using the APRR2_INDEL primers, comparing its amplicon size with those of founders F and G (Figure 4c, Table S1). While founders F and G exhibited an amplicon size of 5,842 bp, the 67/3 accession amplified a shorter fragment of only 566 bp, which was in agreement with the in-silico prediction.

The large deletion present in the 67/3 and HQ-1315 accessions could be responsible for the absence of FC in these genotypes, since they did not present the same high-effect variants found on founders F and G. By analysing the full-length amino acid sequence of *SmAPRR2* in the NCBI conserved domain server (https://www.ncbi.nlm.nih.gov/Structure/cdd/wrpsb.cgi), the 132 truncated residues at the C-terminal region caused the loss of a functional domain (184 amino acids) that corresponded to a golden-2 like transcription factor domain (Figure 5a).

**Figure 5.**
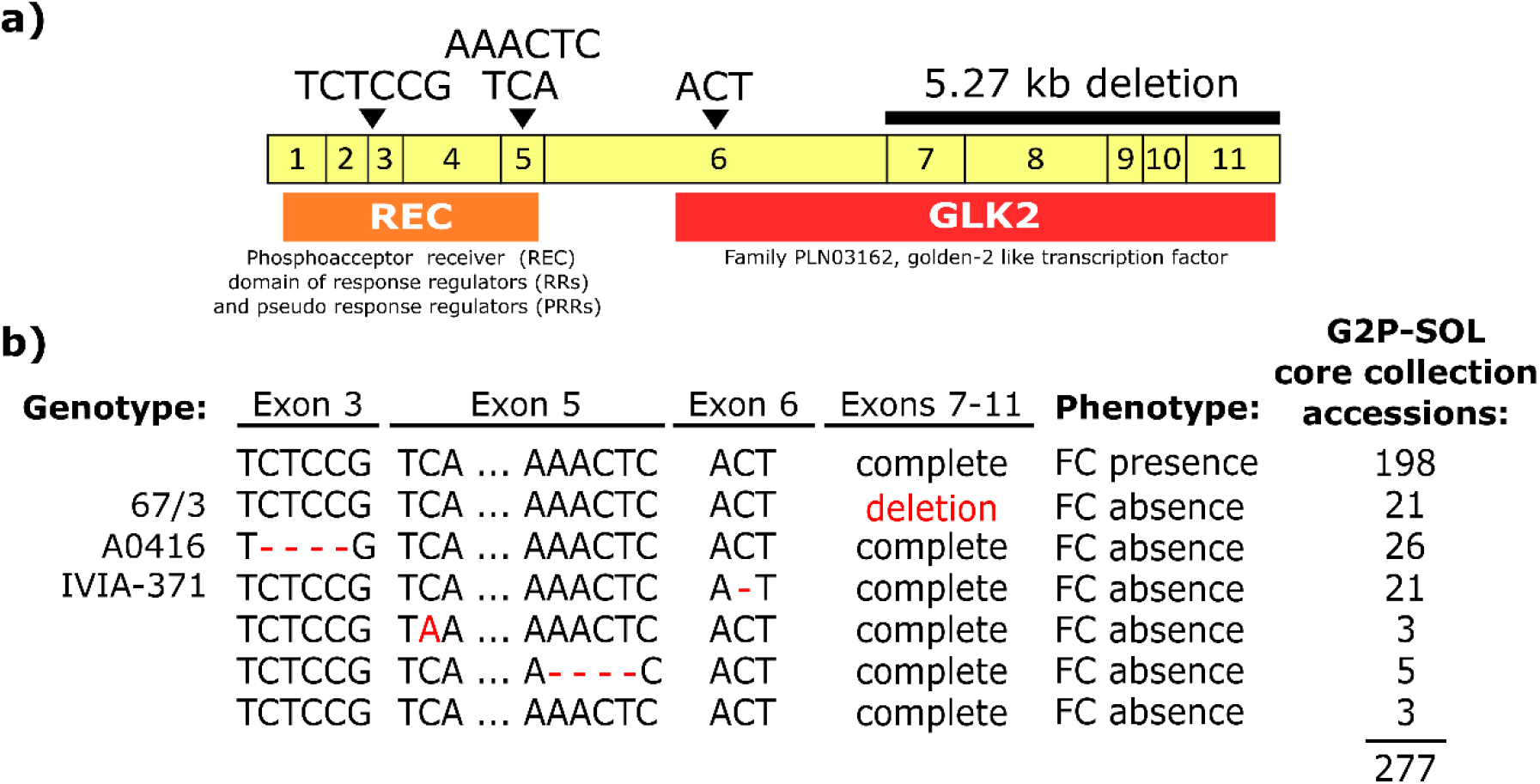
(a) Conservative domain analysis of *SmAPRR2* full-length amino acid gene sequence, including the coding DNA sequence (CDS) indicating each of the exons as yellow boxes, the position of the different variants identified as causative for fruit chlorophyll absence indicated with black arrowheads, the large 5.27 kb deletion and the conserved domains REC and GLK2 as orange and red boxes, respectively. (b) Gene sequence variants identified from the assembly and alignment of 277 accessions from the eggplant G2P-SOL germplasm core collection indicating the genotype, phenotype and the number of accessions carrying each variant.

To investigate the *APRR2* gene structure in relevant species in the Solanaceae and Cucurbitaceae families, a phylogenetic analysis was performed revealing orthology among *APRR2-like* proteins within families, although no clear relationship was found between the Cucurbitaceae and Solanaceae genes (Figure 6 and Figure S2).

**Figure 6.**
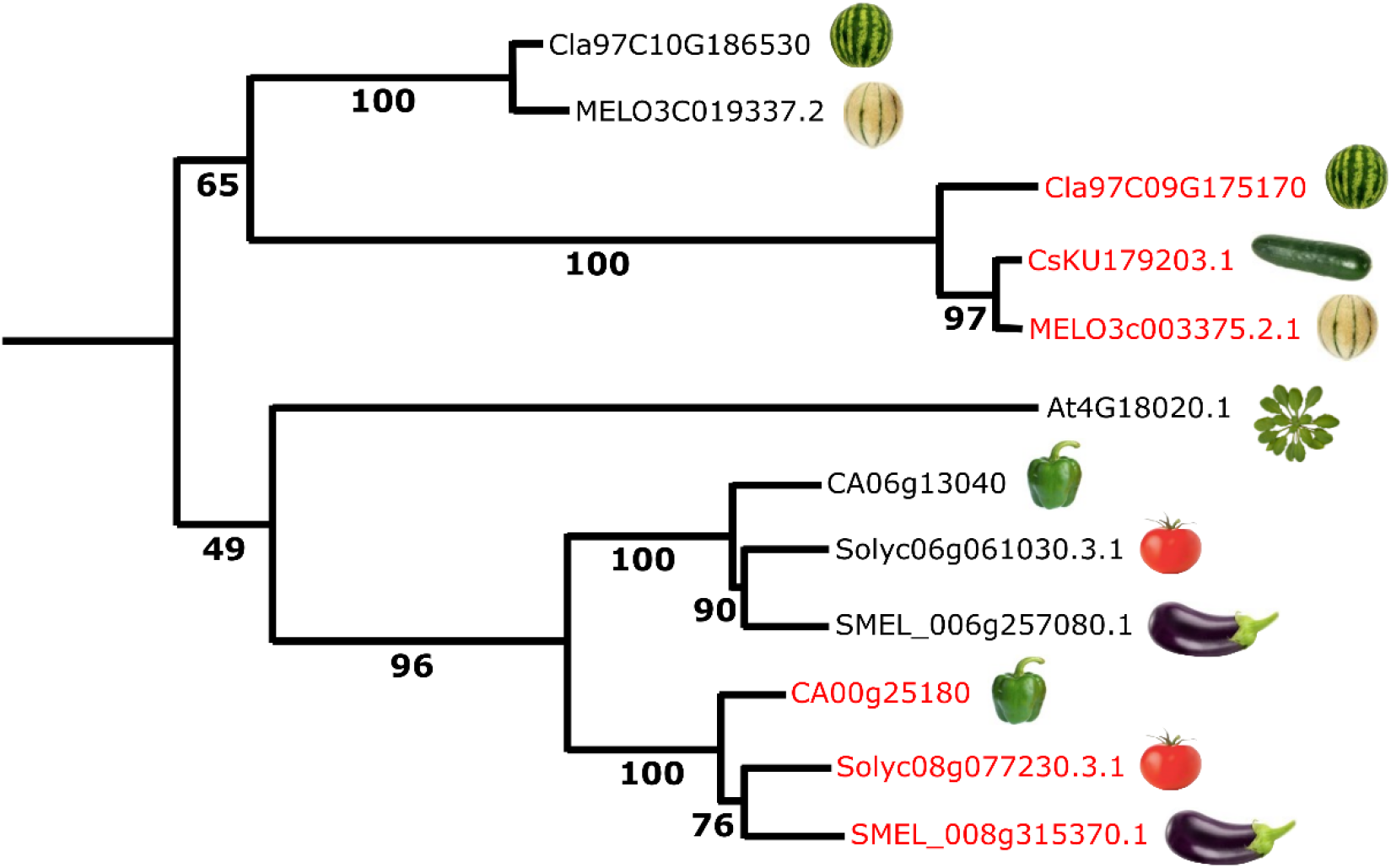
Maximum likelihood dendrogram of *APRR2-like* proteins from *Arabidopsis thaliana*, Cucurbitaceae and Solanaceae species. The red-colored proteins control fruit chlorophyll pigmentation (Pan et al., 2013; Liu et al., 2016; Oren et al., 2019; Jeong et al., 2020; this paper). Cla: *Citrullus lanatus* (watermelon); MELO: *Cucumis melo* (melon); Cs: *Cucumis sativus* (cucumber); At: *Arabidopsis thaliana*; CA: *Capsicum annuum* (bell pepper); Solyc: *Solanum lycopersicum* (tomato); SMEL: *S. melongena* (eggplant). Gene/IDs are from cucurbitgenomics.org (Cucurbitaceae), Solgenomics.net (Solanaceae), Arabidopsis.org (Arabidopsis) and NCBI (cucumber).

### Gene validation in a germplasm collection

The *SmAPRR2* gene sequence was also analysed in a highly diverse germplasm population to validate the hypothesis of *SmAPRR2* as the best candidate gene for FC biosynthesis. The assembly and alignment of 277 accessions from the eggplant G2P-SOL germplasm core collection allowed their classification according to the candidate variants causative for FC absence (Figure 5b, Table S2). Among the 79 accessions without FC, 26.58% (21 of 79) presented the 5.27 kb deletion, the same as the 67/3 and HQ-1315 reference genomes, 32.91% (26 of 79) presented the A0416 (S3MEGGIC population founder F) deletion, and 26.58% (21 of 79) presented the IVIA-371 (S3MEGGIC population founder G) deletion. The remaining 11 accessions exhibited two new high impact variants: three presented a C→A SNP in the 104,011,141 bp position on exon 5, resulting in a premature stop codon, while 5 presented a small A(AACT)C deletion in the 104,011,149 bp position on exon 5, resulting in a frameshift mutation. The remaining 3 accessions without FC had none of the above variants or any apparent high-effect mutation in the sequence *SmAPRR2*. This discrepancy might be caused by phenotyping errors. As expected, for the 198 accessions with FC none of the five variants were observed in the *SmAPRR2* gene sequence.

Furthermore, we checked if there was an association between the geographical origins of the G2P-SOL germplasm core collection accessions and the different variants identified in the *SmAPRR2* gene (Figure 7). It was observed that the 5.27 kb deletion predominated in accessions from Italy (nine of 21, 42.86%), followed by Philippines (four of 21, 19.05%), while the rest of the accessions where mainly distributed among France, China, and India (two of 21 each, 9.52%). The S3MEGGIC founder F (A0416) indel was mainly observed in Thailand and Laos, where 73.08% (19 of 26) and 15.38% (four of 26) of the accessions originated, respectively. Finally, the S3MEGGIC founder G (IVIA-371) indel was distributed mainly between Turkey and Spain with 33.33% (seven of 21) of the accessions having this mutation present in each country, followed by India with a 19.05% (four of 21).

**Figure 7.**
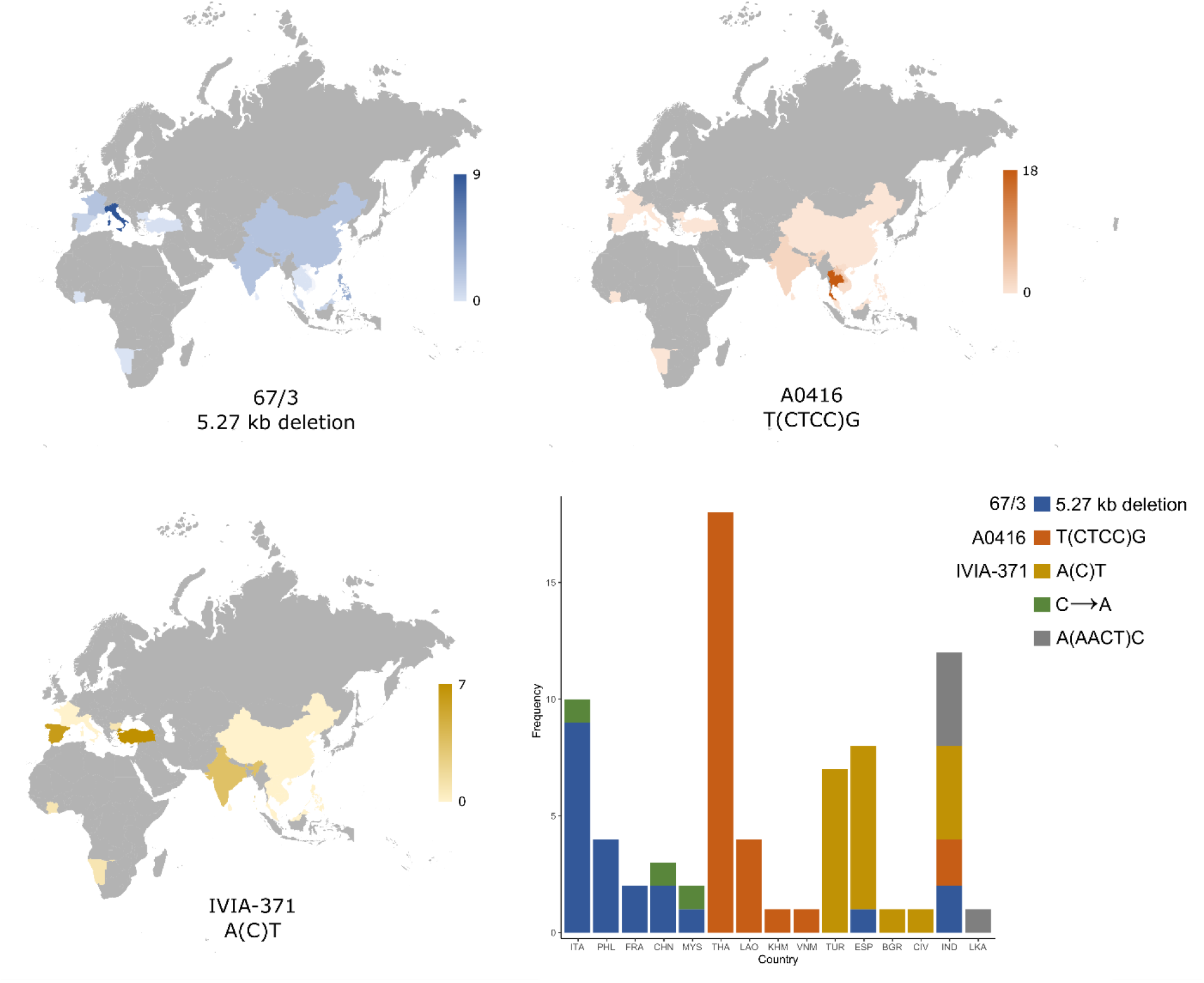
Distribution of the three most abundant high effect variants identified in the *SmAPRR2* gene sequence according to the country of origin of the 277 accessions from the eggplant G2P-SOL germplasm core collection. The numbers on the scale next to each map indicate the numbers of accessions from each country carrying the mutation. Bottom right, a stacked column chart comprising the frequency of the different *SmAPRR2* gene variants across the countries of origin: Italy (ITA), Philippines (PHL), France (FRA), China (CHN), Malaysia (MYS), Thailand (THA), Laos (LAO), Cambodia (KHM), Vietnam (VNM), Turkey (TUR), Spain (ESP), Bulgaria (BGR), Ivory Coast (CIV), India (IND), and Sri Lanka (LKA).

In addition, each high impact variation seemed to be mostly associated with a specific phenotype. The presence of homogeneous fruit anthocyanins with no pigmentation under the calyx was observed for the 5.27 kb deletion in 66.67% (14 of 21) of cases, while completely white fruits or the presence of green netting in a white background was found for the S3MEGGIC founder F (A0416) variant in 88.46% (23 of 26) of cases, and the presence of stripped distribution of anthocyanins on a white background for the S3MEGGIC founder G (IVIA-371) indel was found in 66.67% (14 of 21) of cases. Regarding the two new high impact variants identified, the low number of accessions did not allow to identify a region of predominant distribution, although the A(AACT)C deletion was exclusively located in India and Sri Lanka.

## Discussion

Among fruit quality traits, peel colour is of primary importance because pigments that confer colour are not only associated with visual preferences, but also with nutritional, health, and flavour values (Brand et al., 2014). Eggplant varieties and landraces display different colours at commercial maturity, resulting from a combination of different patterns of presence or absence of anthocyanins and chlorophylls. Although chlorophylls in the peel of fleshy fruits might have a considerable effect on the quality composition of eggplant, as has been demonstrated for tomato (Li et al., 2018b), responsible genes have not been elucidated yet.

To dissect this important trait for eggplant genetics and breeding, in this study we have combined classical forward genetics techniques with advanced association and sequencing analyses, using both experimental populations and germplasm materials to identify and validate a candidate gene controlling the synthesis of fruit peel chlorophyll. Up to now, the dissection of complex traits in eggplant has been usually performed using germplasm panels (Portis et al., 2015) or temporary mapping populations, such as F_2_ and BC_1_ (Barchi et al., 2012; Miyatake et al., 2016; Pandiyaraj et al., 2019; Bhanushree et al., 2020). These populations have the advantage of being easy and fast to develop but are not as powerful as the immortal ones to identify strong associations, candidate genes or causative SNPs, and accumulate fewer genetic recombination events, limiting the resolution for QTL detection (Arrones et al., 2020). In eggplant, this has led to the identification of very few candidate genes controlling traits of interest, lagging it behind other major crops where multiple bi-parental and multi-parent experimental populations have been developed and available for many years already (Gramazio et al., 2018). Luckily, in the last few years, new advanced populations have been developed in eggplant, including recombinant inbred lines (RILs) (Lebeau et al., 2013; Mishra et al., 2020; Toppino et al., 2020; Sulli et al., 2021), and one set of introgression lines (ILs) (Gramazio et al., 2017). These populations have allowed the identification of many and highly relevant QTLs (Mangino et al., 2020; 2021; Toppino et al., 2020; Rosa-Martínez et al., 2022). Eventually, the first eggplant multi-parent advanced generation inter-cross (MAGIC) population has also been developed, allowing the identification of candidate genes for anthocyanin pigmentation in eggplant fruits with greater accuracy and statistical robustness (Mangino et al., 2022).

In this study, this new MAGIC population, called S3MEGGIC, allowed the identification by GWAS analysis of a strong association (LOD = 34.88) for fruit chlorophyll in the peel and the identification of several potential candidate genes beneath or close to the association peak. Furthermore, the whole-genome resequencing of the founders coupled with the high-throughput genotyping of the 420 S3MEGGIC individuals through the identification of “high-impact” variants and haplotype predictions provided strong evidence that ARABIDOPSIS PSEUDO RESPONSE REGULATOR2 *(APRR2)* is the candidate gene for controlling this important trait. In tomato, the overexpression of *SlAPRR2* in transgenic plants resulted in an increased plastid number, area, and pigment content, enhancing chlorophyll levels in immature unripe fruits (Pan et al., 2013). Through the development of two F_2_ and one BC_1_ mapping populations in cucumber, *CsAPRR2* was identified as responsible for the green colour of immature fruits (Jiao et al., 2017). In the case of melon and watermelon, the development of different bi-parental segregating populations and a GWAS panel allowed the association of *CmAPRR2* and *ClAPRR2* as causative genes regulating fruit rind chlorophyll accumulation (Oren et al., 2019). Similarly, in pepper, *CaAPRR2* was found to be strongly associated with chlorophyll pigment accumulation in immature fruit tissues through the analysis of an F_2_ population and confirmed through virus-induced gene silencing (VIGS) (Jeong et al., 2020). Null mutations on *APRR2* have been related in some way to alterations in abscisic acid (ABA) signalling related to plastid development causing reduced chloroplast density and chlorophyll content (Pan et al., 2013; Oren et al., 2019). Here, we present the first evidence of APRR2 as the main actor in chlorophyll presence in the peel of eggplant fruits. Previous studies focused on the eggplant green ring, observed in the flesh next to the skin in an F_2_ population, identified a major QTL on chromosome 8 linked to the marker *35002_PstI_L402* (Portis et al., 2014). Although these authors identified a member of the ferredoxin gene family as the best candidate gene, the marker position is only 623 kb away from the *SmAPRR2* gene.

Thus, *APRR2* homologs have been shown to control fruit chlorophyll content in six different species: three Cucurbitaceae (melon, watermelon and cucumber) and three Solanaceae (tomato, pepper and eggplant). It is noteworthy that the three Cucurbitaceae genes are orthologs of each other, as are the three Solanaceae ones. In contrast, no clear evolutionary relation could be found between the Cucurbitaceae and Solanaceae genes. Thus, the most likely scenario is that the specialization of these *APRR2-like* genes in controlling fruit chlorophyll content occurred early during the evolution of Cucurbitaceae and Solanaceae, and that the two events were independent from each other.

We also reported that the *SmAPRR2* gene sequence for the 67/3 and HQ-1315 eggplant reference genomes, both without fruit chlorophyll, presented a large deletion event of 5.27 kb, disrupting the chlorophyll synthesis in the fruit peel. However, a larger gene size of 4.794 bp with 11 exons was found in the Nakate-Shinkuro and GUIQIE-1 eggplant reference genomes, both with FC (Hirakawa et al., 2014; Li et al., 2021). The availability of different reference genomes contrasting for several traits, like in this study for fruit chlorophyll, fosters their dissection and highlights the need for developing newer, more phenotypically diverse, and accurate reference genomes (Li et al., 2021). The Nakate-Shinkuro and GUIQIE-1 *SmAPRR2* gene structure is largely in agreement with the ones reported in other species, such as the tomato gene *Solyc08g077230* with a 5,831 bp gene size and 11 exons, the cucumber *Csa3G904140.3* with 5,878 bp and 12 exons, the melon *Melo3C003375* with approximately 6,000 bp and 12 exons, the watermelon *CICG09G012330* with 8,561 bp and 11 exons, or the pepper *Capana01g000809* with 5,588 bp and 11 exons (Pan et al., 2013; Jiao et al., 2017; Oren et al., 2019; Jeong et al., 2020). The deletion found in 67/3 and HQ-1315 was associated with 132 residues causing the loss of a functional domain, the golden-2 like transcription factor. A similar deletion in the cucumber *CsAPRR2* gene, encoded for a truncated 101-amino acid protein, was suggested to control the white immature skin fruit colour (Liu et al., 2016). A subcellular localization analysis revealed that the loss of the *CsAPRR2* golden-2 like transcription factor, which acts as a nuclear localization signal domain, caused the impossibility for the protein to enter the nucleus and perform its function (Jiao et al., 2017).

The availability of a large number of accessions, which included a broad range of eggplant genetic diversity, of the eggplant G2P-SOL germplasm core collection made possible the in-silico validation of the hypothesis of *SmAPRR2* as the best candidate gene for fruit peel chlorophyll. Due to the high recalcitrance of in vitro regeneration after *Agrobacterium*-mediated transformation of eggplant (García-Fortea et al., 2020; Mir et al., 2021), routinely reverse genetics gene validation techniques, such as over-expression or knock-out, are still the major bottleneck in this crop. Therefore, the screening of large germplasm collection remains the best choice to indirectly validate gene functions. In fact, despite the explosion of CRISPR/Cas studies in many crops, the first and only eggplant CRISPR plant so far has been obtained by the authors of this study (Maioli et al., 2020), reflecting the inevitable need for alternative indirect validations until new protocols overcome the recalcitrance.

Combining all this information, we highlighted that the two small high-impact variants identified in the S3MEGGIC population, the large deletion of the 67/3 and HQ-1315 reference genomes plus the two high-impact disruptive variants identified in the core collection are responsible for the lack of chlorophyll in the fruit. These five disruptive mutations indicate that the lack of chlorophyll in the eggplant fruit peel has arisen and been selected independently several times, suggesting a prominent role of this trait in eggplant domestication and diversification. Furthermore, we observed that mutated *SmAPRR2* gene variants displayed a different and specific geographical distribution, being also associated with a specific phenotype. This distribution might be related to the domestication flows of eggplant together with local preferences. It has been suggested that two main different domestication events occurred in eggplant, one in India and one in China, since both regions have a high diversity of landraces and populations of putatively wild eggplants (Meyer et al., 2012). Here, we observed that India concentrated the greatest diversity for high impact gene variants for the *SmAPRR2*, including accessions with almost all the identified gene variants. Meyer et al. (2012) also proposed that landraces originated from India would have spread west to western Europe, possibly carried by Arab traders, while landraces arising in China spread to the northeast and southeast Asia. This hypothesis is in agreement with the high frequencies observed of the 5.27 kb deletion present in 67/3 and HQ-1315 and the indel present in the S3MEGGIC founder G (IVIA-371) in Europe, where purple eggplants are popular, and the indel of the S3MEGGIC founder F (A0416) to southeast Asia, where white eggplants are most consumed. This may also suggest that the mutations causing the lack of chlorophyll in gene *SmAPRR2* predate the spread of eggplant to other regions outside its region of origin. In this respect, individuals with white fruits from wild populations of eggplant have been described in India and in other parts of Southeast Asia (Davidar et al., 2015; Page et al., 2019).

Although no clear evidence was found in our population, the Golden 2-like MYB (*GLK2*) gene, which was identified in the genome region where a minor GWAS peak was identified on chromosome 4, is described as a widely conserved transcription factor that positively regulates fruit chlorophylls in different species. Some reports suggest a role of GLK transcription factors in regulating chlorophyll levels in *Arabidopsis*, tomato, and pepper (Fitter et al., 2002; Powell et al., 2012; Brand et al., 2014). The potential tomato ortholog is the *u* (uniform ripening) gene affecting shoulder colour in unripe fruits (Powell et al., 2012). In eggplant, a QTL in chromosome 4 has been related to the reticulated pattern of chlorophylls in the fruit, also called fruit chlorophyll netting or variegation (Daunay et al., 2004; Frary et al., 2014). Although it seems to be independent of the uniform distribution of chlorophyll in the eggplant fruit peel, its presence in fruits could lead to phenotyping errors, which might account for the small discrepancies between the genotype and phenotype that we observed during the experimental procedures.

In conclusion, using different eggplant experimental materials, a MAGIC population and a germplasm core collection, combined with advanced association and bioinformatics analyses and classical genetics tools, we found that *SmAPRR2* is a key gene in the eggplant fruit peel chlorophyll biosynthesis. Our finding is also supported by its conserved function in regulating fruit green pigmentation of different vegetable crops. The dissection of the genetics of this important trait will be extremely useful to foster future breeding programs focused not only on specific market demand based on visual preferences, but also on developing new varieties with improved nutritional quality and resilient against the upcoming environmental challenges due to the implication of chlorophylls in stress biology of plants. The identification of the eggplant *SmAPRR2* represents a landmark for eggplant breeding for fruit colour and other related quality traits.

## Experimental procedures

### Plant materials

A total of 420 S3 individuals from the eggplant S3MEGGIC multi-parent advanced generation inter-cross (MAGIC) population developed by Mangino et al. (2022) were used to identify candidate genes associated with the chlorophyll pigmentation in the fruit peel. The S3MEGGIC population, which is the first and so far the only MAGIC population in eggplant, was obtained by inter-crossing one wild *S. incanum* accession and seven *S. melongena* accessions. Two out of the seven S3MEGGIC *S. melongena* founders (A0416 and IVIA-371; founders F and G, respectively) do not have chlorophyll in the fruit peel, resulting in a population of S3 individuals segregating for this trait. In addition, genomic sequences of 277 *S. melongena* accessions, available from the eggplant germplasm core collection established in the framework of the G2P-SOL project (http://www.g2p-sol.eu/G2P-SOL-gateway.html) and phenotyped for fruit peel colour, were interrogated for the candidate gene identified for controlling chlorophyll pigmentation. This core collection includes accessions used for developing the first eggplant pan-genome (Barchi et al., 2021).

### Fruit chlorophyll phenotyping

The presence of chlorophyll distributed all over the peel of the fruit was screened in the 420 S3MEGGIC individuals and 277 core collection accessions using a binary classification (presence/absence). Fruits were phenotyped at the stage of commercial maturity (e.g., when the fruit is still physiologically immature). When the fruit peel had no anthocyanins or they were distributed irregularly, green pigmentation was easily phenotyped with the naked eye. However, when anthocyanins were distributed all over the fruit peel, the presence of chlorophylls was determined by observing the stylar scar on the distal part of the fruit (Daunay et al., 2004).

### S3MEGGIC Genome-Wide Association Study (GWAS) and haplotype diversity

The genotyping data of the 420 S3MEGGIC individuals, assessed by the Single Primer Enrichment Technology (SPET) eggplant probes available (Barchi et al., 2019a), was retrieved from Mangino et al. (2022). Combining both phenotypic and genotypic data, a Genome-Wide Association Study (GWAS) was performed using the aSSociation, Evolution and Linkage (TASSEL) software (ver. 5.0, Bradbury et al., 2007). For the association study, mixed linear model (MLM) analyses were conducted. The multiple testing was corrected with the Bonferroni and the false discovery rate (FDR) methods (Holm, 1979; Benjamini and Hochberg, 1995) at the significance level of 0.05 (Thissen et al., 2002). The R qqman (Turner, 2014) and LDBlockShow (Dong et al., 2021) packages were used for the Manhattan plots visualization and linkage disequilibrium (LD) determination for haplotype block structure plotting, respectively. LD correlation coefficient (r^*2*^) was used for the pattern of pairwise LD between SNPs measurement, considering haplotype blocks for r^*2*^ values greater than 0.5 and supported by the solid spine of LD method (Gabriel et al., 2002; Barret et al., 2005). Candidate genes found in the most significant regions were first retrieved from the 67/3 eggplant reference genome (ver. 3) (Barchi et al., 2019b). Candidate gene allelic variants were assessed by SnpEff software prediction (ver. 4.2, Cingolani et al., 2012) on the eight S3MEGGIC founders’ resequencing data (Gramazio et al., 2019). Integrative Genomics Viewer (IGV) tool was used for the visual exploration of founder genome sequences to validate SnpEff results (Robinson et al., 2020). Founder haplotypes were estimated for the regions with significant associations using the SNPs with the highest LODs. In addition, a comparative analysis of founder haplotype diversity across the 420 S3MEGGIC individuals was performed by combining genotypic and phenotypic data.

### SmAPRR2 sequence analysis

Allelic variants of the eggplant ARABIDOPSIS PSEUDO RESPONSE REGULATOR2 *(SmAPRR2*) gene were interrogated in the S3MEGGIC population. Of the total 420 S3 individuals, 20 individuals presenting fruit peel chlorophyll (including all combinations of founder haplotypes with chlorophyll presence) and 60 without fruit chlorophyll (30 associated with the A0416 founder haplotype and 30 with the IVIA-371 founder haplotype) were selected to confirm the involvement of the high-impact variants in *SmAPRR2* identified in the resequencing data of the founders. Genomic DNA of the selected individuals was extracted using the SILEX extraction method (Vilanova et al., 2020) and its quality and integrity were checked by agarose electrophoresis and Nanodrop ND-1000 spectrophotometer ratios 260/280 and 260/230 (Thermo Fisher Scientific, Waltham, MA, United States). DNA concentration was estimated with Qubit 2.0 Fluorometer (Thermo Fisher Scientific, Waltham, MA, United States) and adjusted to 50 ng/μL for PCR amplification. In order to genotype the *SmAPRR2* candidate allelic variants in the selected S3MEGGIC individuals, specific primers were designed based on the “67/3” reference genome sequence and founders’ resequencing data (Table S1) and the resulting amplifications were visualized by 1% agarose gel electrophoresis. For long amplification fragments (>5 kb), TaKaRa LA Taq® DNA Polymerase (Takara Bio Inc., Shiga, Japan), which is optimized for long-range PCR up to >15 kb fragments, was used. PCR amplicons were purified with ammonium acetate and sequenced by Sanger sequencing (DNA Sequencing Service, IBMCP-UPV, Valencia, Spain).

To compare the *SmAPRR2* gene structure, the nucleotide sequence was retrieved by a BLASTx search (e-value cut-off of 1e^−5^) against different eggplant reference genomes showing fruit peel chlorophyll, such as the Nakate-Shinkuro and the GUIQIE-1 (Hirakawa et al., 2014; Li et al., 2021), and others showing absence of fruit peel chlorophyll, such as the 67/3 (ver. 3) and the HQ-1315 (Barchi et al., 2019b; Wei et al., 2020). A conservative domain analysis was performed by assessing the NCBI conserved domain server (https://www.ncbi.nlm.nih.gov/Structure/cdd/wrpsb.cgi). To gain insight into the relationship of *APRR2-like* proteins in different species, a phylogenetic analysis was also performed using the predicted protein sequences from *Arabidopsis thaliana*, Cucurbitaceae and Solanaceae families. Protein alignments were performed using MEGA 11.0.10 software (http://megasoftware.net/) and the dendrogram was constructed using the IQtree web server (http://iqtree.cibiv.univie.ac.at/) via maximum likelihood method with default settings.

### Germplasm collection validation

To validate the *SmAPRR2* allelic forms, we examined the GUIQIE-1 eggplant reference genome (Li et al., 2021), since GUIQIE-1 fruits have fruit peel chlorophyll and its assembly is much more complete than the Nakate-Shinkuro reference genome (Hirakawa et al., 2014). Furthermore, whole-genome resequencing data of the 277 eggplant G2P-SOL germplasm core collection were interrogated (unpublished data). After trimming with SOAPnuke software (Chen et al., 2018) with filter parameters: “-l 20 -q 0.5 -n 0.03 -A0.28”, reads were aligned against GUIQIE-1 genome using the Burrows-Wheeler Aligner (BWA) program (v. 0.7.17– r1188, Li and Durbin, 2009) and the ‘mem’ command with default parameters. Subsequently, reads mapping in the genomic region of *SmAPRR2* were extracted with samtools and de novo assembled using Megahit (v.1.2.9, Li et al., 2015) with default parameters. The sequencing raw data of the newly sequenced accessions are available at NCBI SRA (BioProject ID PRJNA837769).

A multiple sequence alignment of the assembled sequences was then performed using the MAFFT program (ver. 7, Katoh et al., 2008) and the results were visualized in the Jalview alignment editor (ver. 2, Waterhouse et al., 2009). The resulting data were used to associate the different variants identified in the *SmAPRR2* gene for the G2P-SOL core collection and the origin of the accessions.

## Author Contributions

JP, SV and PG conceived the idea and supervised the study. AA, GM, DA, MP and PG performed the field trials. AA, GM, SV and PG performed the analysis of the S3MEGGIC population and the *APRR2* gene structure. LB, EP and GG performed the analyses of the G2P-SOL core collection and the phylogenetic analysis. AA and PG prepared a first draft of the manuscript and the rest of the authors reviewed and edited the manuscript. All authors have read and agreed to the published version of the manuscript.

## Acknowledgments

This work has been funded by grants RTI-2018–094592-B-I00 and PID2021-128148OB-I00 funded by MCIN/AEI/10.13039/501100011033/ and by “ERDF A way of making Europe”, and by European Union’s Horizon 2020 Research and Innovation Programme under Grant Agreement No. 677379 (G2P-SOL project: Linking genetic resources, genomes and phenotypes of Solanaceous crops). AA is grateful to Spanish Ministerio de Ciencia, Innovación y Universidades for a pre-doctoral (FPU18/01742) contract. PG is grateful to Spanish Ministerio de Ciencia e Innovación for a post-doctoral grant (FJC2019-038921-I/AEI/10.13039/501100011033). Funding for open access charge: Universitat Politècnica de València.

## Supporting Information

**Figure S1**. Local Manhattan plot and LD heatmap of the candidate peaks on chromosomes 8 and 4, top and bottom figures, respectively. The red and green horizontal lines represent, respectively, FDR and Bonferroni significance thresholds at p=0.05. In red are represented the points that exceed these thresholds, while in blue are the points that do not. Pairwise LD between SNPs is indicated as r2 values: red indicates a value of 1 and white indicates 0.

**Figure S2**. MEGA alignment of *APRR2-like* protein sequences from relevant species of the Solanaceae and Cucurbitaceae families and *Arabidobsis thaliana*.

**Table S1**. List of primers used in this study.

**Table S2**. Gene sequence variants identified in the 277 accessions from the eggplant G2P-SOL germplasm core collection indicating the G2P-SOL ID, the variant present in each accession and the fruit peel chlorophyll phenotype presence (YES) or absence (NO).

