## Supplementary material for "Mutations in the *SmAPRR2* transcription factor suppressing chlorophyll pigmentation in the eggplant fruit peel are key drivers of a diversified colour palette": Figure S2: Supplementary Figure S2.pdf

```
MELO3C019337.2        ------------------------------------------------------------   0
Cla97C02G036020       ------------------------------------------------------------   0
At4G18020.1           ---------------------------------MVITANDLSKWENFPKGLKVLLLLNGC   27
Cla97C09G175170       ---------------------------------MVCTVDDLQEWKDFPKGLRVLLL----   23
MELO3C003375.2.1      ---------------------------------MVCTADDLQEWKDFPKGLRVLLL----   23
CsAMJ39435.1          ---------------------------------MVCTADDLQEWKDFPKGLRVLLL----   23
MELO3C013874.2.1      ---------------------------------MVCTANDLHGWKDFPKGLRVLLL----   23
Cla97C10G186530       ---------------------------------MVCTANDLHGWKDFPKGLRVLLL----   23
CA06g13040            ---------------------------------MVCTENDLLGWKDFPKGLRVLLL----   23
SMEL_006g257080.1     ---------------------------------MVCTENELLEWKDFPKGLKVLLL----   23
Solyc06g061030.3.1    ---------------------------------MVCTENELLEWKDFPKGLKVLLL----   23
SMEL_008g315370.1     ---------------------------------MICIEDELLGWKDFPKGLKVLLL----   23
Solyc08g077230.3.1    ---------------------------------MICIENELLGWKDFPKGLKVLLL----   23
CA00g25180            MFAGRDGFVAVVVLMIGISGVVNGPGAGCNGREMICIEDELLGWKDFPKGLKVLLL----   56

MELO3C019337.2        ------------------------------------------------------------   0
Cla97C02G036020       ------------------------------------------------------------   0
At4G18020.1           DSDGDGSSAAETRSELESMDYIVTTFTDETEALSAVVKNPESFHIAIVEV-NMSAESESF   86
Cla97C09G175170       --DRDSRSASEIRSKLEEMEYVVYSCSDEKEALSAILNTPGNFHVAILEV-CAKNHDESF   80
MELO3C003375.2.1      --DRDSCSATEIRSKLEEMEYVVYSCTDEKEALSAILNTPGNFHVAILE-----------   70
CsAMJ39435.1          --DRDSFSATEIRSKLEEMEYVVYSCTDEKEASSAILNTPGNFHVAILEV-CARNYDESF   80
MELO3C013874.2.1      --DGDTSSAAEIKTKLEEMEYVVVSYCNENDALSAISSKPETFHVAIVEV-TTSNHEGNF   80
Cla97C10G186530       --DGDSSSAAEIKTKLEEMEYVVVTYCNENDALSAISSKPETFHVAIVEV-TTSNHNGNF   80
CA06g13040            --DKDSNSASDMRSRLEEMEYIVYAFCNETEALSAISSKSEVFHVAIVEV-SAGNSDGGL   80
SMEL_006g257080.1     --DKDCSSASQMRSRLQEMDYIVHTFCNENEALSAISSKSEVFHVAIVEV-SDGNSDGEL   80
Solyc06g061030.3.1    --DTDSNFASQMRSRLQQMDYIVYTFCNENEALSAISSKSEVFHVAIVEV-SAGNSDGGF   80
SMEL_008g315370.1     --DEDSNSAAEMRSRLEKMDYIVYSFCNESEALTAISSKSEGFHVAIVEV-SEGNSDGVL   80
Solyc08g077230.3.1    --DEDSNSAAEMKSRLEKMDYIVYSFCNESEALTAISSKSEGFHVAIVEV-SAGNSDGVL   80
CA00g25180            --DEDSNSAAEMKSRLEKMDYIVYTFCNESEALSAISSKSEGFHVAIVEVVSAGDNDGVL   114

MELO3C019337.2        ------------------------------------------------------------   0
Cla97C02G036020       ------------------------------------------------------------   0
At4G18020.1           KFLEAAKDVLPTI-----------------------------------------------   99
Cla97C09G175170       KLLGTSKD-LPII-----------------------------------------------   92
MELO3C003375.2.1      ------------------------------------------------------------   70
CsAMJ39435.1          KLLGASKD-LPII-----------------------------------------------   92
MELO3C013874.2.1      KFLEAAKD-LPTI-----------------------------------------------   92
Cla97C10G186530       KFLEAAKD-LPTI-----------------------------------------------   92
CA06g13040            KFLEGAKD-LPTI-----------------------------------------------   92
SMEL_006g257080.1     KFLESAKD-LPTI-----------------------------------------------   92
Solyc06g061030.3.1    KFLESAKD-LPTI-----------------------------------------------   92
SMEL_008g315370.1     RFLESAKD-LPTIKQFGGHYRMESLCGRYIMETPELGIVCSSSNVINAKTYLRELHLLCA   139
Solyc08g077230.3.1    RFLESAKD-LPTI-----------------------------------------------   92
CA00g25180            QFLESAKN-LPTI-----------------------------------------------   126

MELO3C019337.2        ------------------------------------------------------------   0
Cla97C02G036020       ------------------------------------------------------------   0
At4G18020.1           MISTDHCITTTMKCIALGAVEFLQKPLSPEKLKNIWQHVVHKAFNDGGSNVSISLKPVKE   159
Cla97C09G175170       MTSDVHCLSTMMMCIALGAVEFLLKPLSEDKLRNIWQHVIHKAFSNT-------SKPDED   145
MELO3C003375.2.1      ----------------LGAVEFLLKPLSEDKLRNIWQHVIHKAYSNT-------SKPGEE   107
CsAMJ39435.1          MTSDVHCLSTMMKCIALGAVEFLLKPLSEDKLRNIWQHVIHKAYSNS-------SKPDED   145
MELO3C013874.2.1      MISNIHCLSTMMKCIALGAMEFLQKPLSDDKLRNIWQHVVHKAFNAGGSAVPNSLKPIKE   152
Cla97C10G186530       MISNIHCLSTMMKCIALGAIEFLQKPLSEDKLRNIWQHVVHKAFNAGGSPVPNSLKPIKE   152
CA06g13040            MVSNIHSISTMMKCIALGAVEFLQKPLSDDKLRNIWQHVVHKAFHSGGKNVAESLKPVKE   152
SMEL_006g257080.1     MVSNIHSISTMMKCIALGAVEFLQKPLSDDKLRNIWQHVVHKAFHSGGKNVSESLKPVKE   152
Solyc06g061030.3.1    MVSDIHSINIMMKCIALGAVEFLQKPLSDDKLRNIWQHVVHKAFHSGGKSVSESLKPVKE   152
SMEL_008g315370.1     VTSNIHSLSTMMKCIALGAVEFLQKPLSDDKLKNIWQHVVHKAFNS-RKDVSRSLDPVKE   198
```

```
Solyc08g077230.3.1      MTSNIHSLSTMMKCIALGAVEFLQKPLSDDKLKNIWQHVVHKAFNT-RKDVSKSLEPVKD    151
CA00g25180              MTSNIHSLSTMMKCIALGAVEFLQKPLSDDKLKNIWQHVVHKAFNT-RKDVSGPLEPVKE    185


MELO3C019337.2          ----MLALSPIRSGNKDEKQGEMERFSI----------GGDDFP-----DFDDDTNLLDS    41
Cla97C02G036020         ----MLALSPIRSGNKDEKQGEMERFSI----------GGDDFP-----DFDDDTNLLDS    41
At4G18020.1             SVVSMLHLETDMTI-------------------------EEKDPAPSTPQLKQDSRLL-D    193
Cla97C09G175170         SIASLMQLQLENEDK-NGVPEDMEILSWIQDIVWEQPEGSGNS----------------Q    188
MELO3C003375.2.1        SVASLMQLQLENEDK-NGVPEDMEILSWIQDIVWEQPEGSDDKS-----QLNLGASRQGS    161
CsAMJ39435.1            SVASLMQFQLQNEDK-NGVPEDMEILSWIQDIVWEQPEGSDDRS-----QLNLGASRQAS    199
MELO3C013874.2.1        SVVSMLHLELENSENENQVEKKLEILSGDDDNNHELLEGSDKYPAPSTPQQKHGMRLVDD    212
Cla97C10G186530         SVASMLHLELENSENENQIQKELEISSRNDDNDHELLEGSDKYPAPSTPQQKHGMRLVDD    212
CA06g13040              SLLSMLELQPVKREADSENSNEAEPLTSVLENQKESPNCCDKYPAPSTPQHKQGVRSVDD    212
SMEL_006g257080.1       SLLSLLELQQVKRE----DTNEAEPLTSVEENQKESPPCCDKYPAPSTPQQKQGVRSVDD    208
Solyc06g061030.3.1      SLLSILELQPVKHEADNENTNEAEPLISVVENQKASSSCCDKYPAPSTPQQKQGVRSVDD    212
SMEL_008g315370.1       SLLSMLQLKPAKDEADDKNSNRIEPLTAIAESNTEQLSGCDKYPAPSTPQLKQGVRSVDD    258
Solyc08g077230.3.1      SVLSMLQLQLEMGEADDKSSNGTEPPTAVAESNTEQSSGCDKYPAPSTPQLKQGVRSVDD    211
CA00g25180              SLLSMLQLQPEKGEPDDKSSNGTEPLIAVADNNTEQSSGCDKYPAPSTPQLKQGVRSVDD    245
                            :: :.                                .                 .

MELO3C019337.2          INFDDLFVGINDGDVLPDLEMDPELLAEFSVSGGEESEVNASVSLEKFDDNTLKIIGNKD    101
Cla97C02G036020         INFDDLFVGINDGDVLPDLEMDPELLAEFSVSGGEESEVNASVCLEKFDDNTKAID----    97
At4G18020.1             GDCQENINFSMENVNS---STEKDNMEDHQD-IGESKSVD-TTNRKLDDD----------    238
Cla97C09G175170         LNLGDQMNCSMETDCR---DKD-----------VQSKFLE-TTSHDLVCEGPLPE-----    228
MELO3C003375.2.1        WESGDQMNCSMETDCK---DKD-----------VQSKFVE-TTSHDLVCEGPIQE-----    201
CsAMJ39435.1            WESGDQMNCSMETDCK---DKD-----------VQSKFVE-TTSHDLICEGPIQE-----    239
MELO3C013874.2.1        GDCQDQLNSSLEKECG---EQD-----------GESKSVE-TTCINSLVEGTSQV-----    252
Cla97C10G186530         GDCHDQLNSSLEKECG---EQD-----------GESKSVE-TTCINSLGEGTSQV-----    252
CA06g13040              GDFQDHTILSNEQDSE---VHE-----------GDTKSVE-TTCCDSVAETSILA-----    252
SMEL_006g257080.1       NDLQDHTILSNEQDSG---VHE-----------GDTKSVE-TTCCGSIAETAVLA-----    248
Solyc06g061030.3.1      VDYQDHTILSNEQDSG---MHE-----------GDTKSVE-TTSCDSVAETTVLA-----    252
SMEL_008g315370.1       GDCHDHTIFSTDQDSG---EHD-----------GDTKSVE-TTYNNSLAENTVQT-----    298
Solyc08g077230.3.1      GDCHDHTIFSTDQDSG---EHD-----------ADTKSVE-TTYNNSLAENNVQT-----    251
CA00g25180              SDCHDHTIFSTDQDNG---EHD-----------GDTKSVE-TTYNNSLAENTVQI-----    285
                         :  :      :         :            : . :: :.      :

MELO3C019337.2          NDDDEDQKDLDSRSSSQ---VVDQEILSKRDDELATPTN---------------------    137
Cla97C02G036020         NKDEDEQKELDSRSCGQGVESIDKEIVSKP-DELATPTN---------------------    135
At4G18020.1             ------------------KVVVKEERGDSEKEEEGET-----------------------    257
Cla97C09G175170         -----------GQPQLSDK---------NKSGV---------------------------    241
MELO3C003375.2.1        -----------GQPQLSDKRVTFIIPLQKKFGV---------------------------    223
CsAMJ39435.1            -----------GQPQLSDK---------KKIGV---------------------------    252
MELO3C013874.2.1        -----------ENSQLPDREAIKEEENSADGCGAASNIDH--------------------    281
Cla97C10G186530         -----------ENSQLPDQEGIKEEENSADGCGAASNIDH--------------------    281
CA06g13040              -----------DSAGRLEVAITKDERDSAADQNMEDPIAT--------------------    281
SMEL_006g257080.1       -----------DSA----RAITKEEHDSAVDQNMEDPIAT--------------------    273
Solyc06g061030.3.1      -----------DSSERLGEAITKEEHYSAADQHMEDPIAT--------------------    281
SMEL_008g315370.1       -----------SPPGQQGERILKEENVSSPHQKMEANIAT--------------------    327
Solyc08g077230.3.1      -----------SPTVQQGDIILKEDNVSSPDLKTETDIAT--------------------    280
CA00g25180              -----------SPPGQQQDIILKEENGSSPHQTMEADIATFSQINDCADNSDGSSPHQKT    334


MELO3C019337.2          ----IIE---ANPLVKDSGDKSIKPQKAS--SSQSKNSQGKRKVKVDWTPELHRRFVQAV    188
Cla97C02G036020         ---INIE---GNSLVKDGGDKIIKPQKCS---SQSKNSQGKRKVKVDWTPELHRRFVQAV    186
At4G18020.1             -GDLISEKTDSVDI-HKKEDETKPINKSSGIKNVSGNKTS--RKKVDWTPELHKKFVQAV    313
Cla97C09G175170         ---------------KSSPLAAEHSIQGSDVNHSAGTK--AKKTKVDWTSELHGKFVQAI    284
MELO3C003375.2.1        ---------------ESDPLAAENSIQGTGVNQSAGSK--AKKTKVDWTPELHRKFVQAV    266
CsAMJ39435.1            ---------------KSDPLAAENSIQGTGVNQSAGSK--AKKTKVDWTPELHRNFVQAV    295
MELO3C013874.2.1        ----------DTHDQYNISSSEKNKSKPCGLSNPCGIKVSRKKLKVDWTPELHRKFVQAV    331
Cla97C10G186530         ----------DTHDRDNISSSEKNKTKPCGVNNPCGTKVSRKKLKVDWTPELHRKFVQAV    331
CA06g13040              -------CSRSNDYPADGSTRSAESNKASGLHSSSGTKANKKKMKVDWTPELHKKFVKAV    334
SMEL_006g257080.1       -------CNDCP---INSSIGSAHRNKASGVHSSSGTKANTKKMKVDWTPELHKKFVKAV    323
Solyc06g061030.3.1      -------CSPSN---DNGSTCSADPNKASGLHSSSGTKANKKKMKVDWTPELHKKFVKAV    331
SMEL_008g315370.1       -------SSQSNDCPDSSISHSAEPSKASGPHSSSGTKSNKKKLKVRWKKWCLV-FIKK-    378
Solyc08g077230.3.1      -------TSRSNDCPDNSIMHSAEPSKASGPHSSNGTKSNRKKIKVDWTPELHKKFVQAV    333
CA00g25180              EADIATTSSQSKDCPDNSISHSAEPSKASGPHSSSGTKSNKKKVKVDWTPELHKKFVQAV    394
                                        .         :          .    : ** *.      *::
```

```
MELO3C019337.2        EQLGVDKAVPSRILELMGIECLTRHNVASHLQ-KYRSHRKHLLAREAEAASWSQRRQMYG   247
Cla97C02G036020       EQLGVDKAVPSRILELMGIECLTRHNVASHLQ-KYRSHRKHLLAREAEAASWSQRRQMYG   245
At4G18020.1           EQLGVDQAIPSRILELMKVGTLTRHNVASHLQ-KFRQHRKNILPKDDHNHRWIQSRENHR   372
Cla97C09G175170       EQIGIDHAIPSKILELMKVEGLTRHNVASHLQ-KYRMQKKHIMQREEN-P-------R-C   334
MELO3C003375.2.1      EQLGIDHAIPSKILELMKVEGLTRHNIASHLQ-KYRMQKKHVMQREEN-TRWSHYP-R-S   322
CsAMJ39435.1          EQLGIDHAIPSKILELMKVEGLTRHNIASHLQ-KYRMQKKHVMQREEN-TRWSHYPTR-S   352
MELO3C013874.2.1      EQLGVNQAIPSRILELMKVEGLTRHNVASHLQ-KYRMHKRHILPKEED-GSWSHSK---D   386
Cla97C10G186530       EQLGVNQAIPSRILELMKVEGLTRHNVASHLQ-KYRMHKRHILPKEED-GSWSHSK---D   386
CA06g13040            EKLGIDQAIPSRILELMKVEGLTRHNIASHLQ-KFRMQRRQILPKEDE-KRWPRPQLR-D   391
SMEL_006g257080.1     EKLGIDQAIPSRILELMKVEGLTRHNIASHLQ-KFRMQRRQILPKEDE-RRWPRPQPR-D   380
Solyc06g061030.3.1    EKIGIDQAIPSRILELMKVEGLTRHNIASHLQ-KFRMQRKQILPKEDE-RRWPRPQPR-D   388
SMEL_008g315370.1     ------------------------------------------------------------   378
Solyc08g077230.3.1    EQLGIDQAIPSRILDLMKVEGLTRHNVASHLQ-KYRMHRKQILPKEVE-RRWPNPQPI-D   390
CA00g25180            EQLGIDQAIPSRILDVMKVEGLTRHNIASHLQQKYRMHRRQILPREVE-RRWPHPQPR-D   452

MELO3C019337.2        GG-GGGGGKREVSSWGAPPTMGFPPMTP-MH--PHFRPLHVWGHPPAMDQSLL-HVWPKH   302
Cla97C02G036020       GAAGGGGGKREVSSWGA-PTMGFPPMTT-MH--PHFRPLHVWGHP-TMDQSLM-HVWPKH   299
At4G18020.1           PN----QRNYNVFQQQHRPVMAYP----------------VWGLPGVYPPGAIPPLWPP-   411
Cla97C09G175170       TM----QTNH------LKPIMAYP-SYHPNCGISVSAVYPTWRQTNGHPANFN----HP-   378
MELO3C003375.2.1      TL----QTNH------LKPIMAYP-SYHPNCGISVSAVYPTWRQTNDRPPNIH--VCGP-   368
CsAMJ39435.1          TL----QTNH------LKPIMAYP-SYHPNCGISVSAVYPTWRQTNDHPPNVH--VWGP-   398
MELO3C013874.2.1      PM----RKN----YYPQRPVMAFPPPYHSNHIMPVAPIYPPWGHMACPSPGVR--MWVP-   435
Cla97C10G186530       PM----KKN----YYPQRPVMAFPPPYHSNHIMPVAPIYPPWGHMACPSPGVQ--MWVP-   435
CA06g13040            SV----QRTY---YYPHKPVMAFP-TYHPNNAPTAGQFYPPWIPPGGYPNGAH--MWGS-   440
SMEL_006g257080.1     LV----QRTY----YPHKPVMAFP-TYHSNHASTAGQFYPAWIPPGGHPNGAH--MWGS-   428
Solyc06g061030.3.1    PV----QRTY----YPHKPVMAFP-THHSNHATTAGQFYPAWIPPGGYPNGAH--MWNS-   436
SMEL_008g315370.1     ------------------------------------------------------------   378
Solyc08g077230.3.1    SV----QRSY----YPHKPIMTFP-QYHSNHVAPGGQFYPAWVTPASYPNGLQ--VWGS-   438
CA00g25180            SV----QRNY----YPHKPVMTFP-PYHSNHVAPAGQCYPAWVPPASYPNGLQ--VWGS-   500

MELO3C019337.2        LPHSPSPPPPPPTPPPSSWPHAAAPPPPPDPSYWHHHHQRVPNGLTSGTPCFPQPIPTTR   362
Cla97C02G036020       LPHSPSPPP-PPTPPPSSWPHAAAPPPPPDPSYWHHHHQRVPNGLTSGTPCFPQPIPTTR   358
At4G18020.1           -------PL-QSIGQPPPWHWKPPY-PTVSGNAW-----GCPVGPPVTGSYITPSNT---   454
Cla97C09G175170       -------PGYRHWPQPGIQPWN-SY-AGVRADAW-----GCPVMLPSHTPYFSYPQH---   421
MELO3C003375.2.1      -------FGYRHWPQPGIQPWN-SY-ARVQADTW-----GCPVMPPSHAPYFSYPQL---   411
CsAMJ39435.1          -------LGYRHWPQPGIQPWN-SY-AGVQADTW-----GCPVMPPSHAPYFSYPQL---   441
MELO3C013874.2.1      -------PGYPPWRPPEIWPWK-SY-PGMHADTW-----GCPVTPLPHSPLSSHPQ----   477
Cla97C10G186530       -------PGYPPWRPPEIWPWK-SY-PGMHADTW-----GCPVTPPPHSPLSSHPQQ---   478
CA06g13040            -------PYYPGWQPPENWHWN-PH-SGLYADVW-----GCPVTPPSLGSCTPYLQ----   482
SMEL_006g257080.1     -------PYYPGWPPPETWHWN-PQ-PGLYADVW-----GCPVTPPSLGSCTPYPQ----   470
Solyc06g061030.3.1    -------PYYHGWQPPETWHWN-PQ-PGLYADVW-----GCPVTPPSLGSCTPYPQ----   478
SMEL_008g315370.1     ------------------------------------------------------------   378
Solyc08g077230.3.1    -------PYYPGWKPAETWHWT-PR-PELHADTW-----GSPIMSPSLGSYPPYPQ----   480
CA00g25180            -------PYYPGWQPAETWHWK-PH-PGLLADTW-----GSPVMPPSFGSYPPYPQ----   542

MELO3C019337.2        FGGASFSVIPPPHPMYKAAEPTTSVGRSPTHPPLDSYPSKESIDSAIGDVLAKPWLPLPL   422
Cla97C02G036020       FGGAGFSVVPPPHPMYKA-EPTATVGRSPTHPPLDSYPSKESIDSAIGDVLAKPWLPLPL   417
At4G18020.1           -TAGGFQYPN-----------GAETGFKIMP---ASQPDEEMLDQVVKEAISKPWLPLPL   499
Cla97C09G175170       -VSA-SHNMH-----------TVNKSYGMPQGLFDLQPDEKVVDKIVKEAMRKPWSPLPL   468
MELO3C003375.2.1      -VSASQHNMH-----------TVNKSYGMPQGLFDLQPDEEVVDKIVKEAMREPWSPLPL   459
CsAMJ39435.1          -VSASQHNMH-----------TVNKSYGMPQGLFDLQPDEEVVDKIVKEAMKKPWSPLPL   489
MELO3C013874.2.1      -HISGFENAD-----------PYDKSYSIAFSPIDLQLADEEIDKVVKEAISKPWLPLPL   525
Cla97C10G186530       -HISRFESTD-----------PYDKSYGIAFSPIDLQLADEEIDKVVKEAISKPWLPLPL   526
CA06g13040            -NAS-----------------GIHNRYGIIQKSVDLHPAEEVIDKVVKEAIHKPSLPLPL   524
SMEL_006g257080.1     -NASRFHRAE-----------GIHNRYSSIEKSVDLHPAEEVIDKVVKEAINKPWLPLPL   518
Solyc06g061030.3.1    -NASGFHRAE-----------GMLNGYSIIQKSVDLHPAEEVIDKVVKEAINKPWLPLPL   526
SMEL_008g315370.1     ------------------------------------------------------------   378
Solyc08g077230.3.1    -NAGV-YRPH-----------GTHNRYSMLEKSFDLHPADEVIDKVVKEAITKPWLPLPL   527
CA00g25180            -NAGM-YQSH-----------GMHNRYSMLEKSFDVHPAEEVIDKVVKEAITKPWLPLPL   589

MELO3C019337.2        GLKPPSLDSVKVELQRQGIPKIPPTCAA--------    450
Cla97C02G036020       GLKPPSLDSVKVELQRQGIPKIPPTCAAA-------    446
```

```
At4G18020.1             GLKPPSAESVLAELTRQGISAVPSSSCLINGSHRLR    535
Cla97C09G175170         GLKPP-TESVLTELSKQGISTVPPR---INGSKPP-    499
MELO3C003375.2.1        GLKPPSTESVLTELSKQGISTVPPQ---IDGSRSP-    491
CsAMJ39435.1            GLKPPSTESVLTELSKKGISTVPPQ---IDGSRSP-    521
MELO3C013874.2.1        GLKPPSTESVLSELSKQGISTVPSH---INGSKVIQ    558
Cla97C10G186530         GLKPPSTESVLSELSRQGISTVPSH---INGSKLLQ    559
CA06g13040              GLKSPSTESVLDALSKQGISAVPSR---INGSRRPH    557
SMEL_006g257080.1       GLKPPSTESVLDALSKQGVSTVPPR---INGSHRPH    551
Solyc06g061030.3.1      GLKPPSTESVLDALSKQGIPAVPPR---HHRSHRPH    559
SMEL_008g315370.1       ------------------------------------    378
Solyc08g077230.3.1      GLKAPSTESVLDELSRQGISTIPSQ---INDSRCRR    560
CA00g25180              GLKPPSMEGVLDELSRQGISTVPPR---INGSRCWR    622
```
