## Supplementary figures and images for "Mutations in the *SmAPRR2* transcription factor suppressing chlorophyll pigmentation in the eggplant fruit peel are key drivers of a diversified colour palette"

### Supplementary Figure S1.png

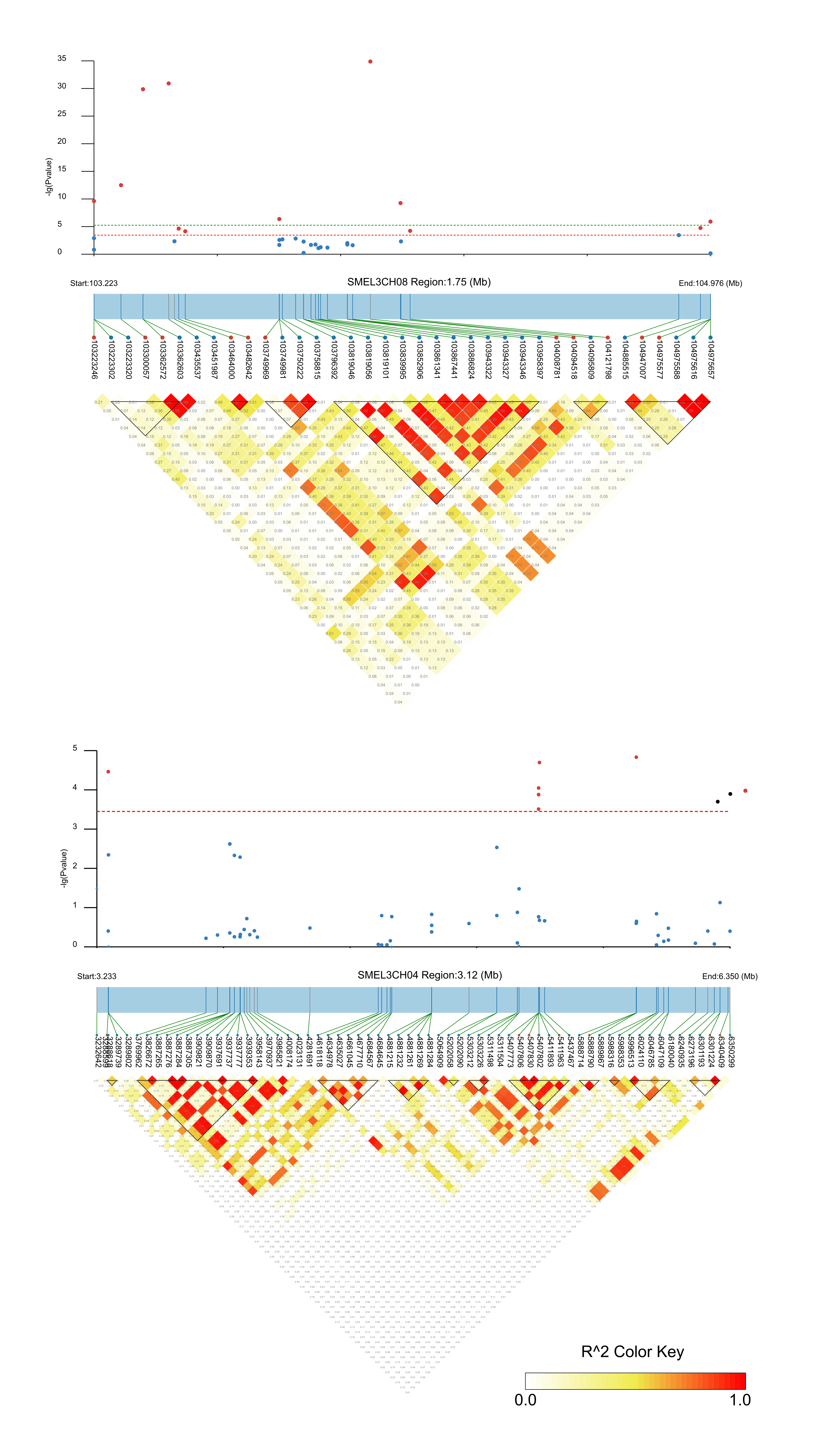
